# DEER of Singly Labelled Proteins to Evaluate Supramolecular Packing of Amyloid Fibrils

**DOI:** 10.1101/2025.11.10.687719

**Authors:** Karen Tsay, Asif Equbal, Yuanxin Li, Tiffany Tsui, Songi Han, Yann Fichou

## Abstract

1

The formation of protein amyloid fibrils in the brain is a hallmark of various neurodegenerative diseases, including Alzheimer diseases and Parkinson diseases. Amyloid fibrils are highly ordered aggregates in which proteins folded in two-dimensional layers stack along one axis to form elongated linear assemblies. The specific conformations adopted by the proteins within each layer of the amyloid correlates with the pathology. Furthermore, their spreading and templating competency relies on strict in register packing of the folded proteins along the fibril growing axis. There is a need for tools to characterize not only the protein fold across the fibril cross section but also the spatial ordering of the proteins stacked along the amyloid fibril axis. We present an approach based on double electron electron resonance spectroscopy (DEER) using singly labelled tau protein assembled in amyloid fibrils that can deliver an apparent dimensionality of the supramolecular organization of tau fibrils. The parameters of the DEER background function can be used to assess the amyloid core location and packing order, and track time-resolved formation of aggregation intermediates. Showcasing the method on tau, we demonstrate that heparin-induced tau fibrils are mispacked while seeded aggregation can template amyloid fibrils with a higher packing order. This study benchmarks a new method that will provide critical structural insights into amyloid assemblies.

DEER-derived dimensionality parameter differentiates between spin-labelled tau protein in solution, in perfectly aligned fibril or in imperfectly aligned fibril (fibril structure edited from PDB 6qjh).

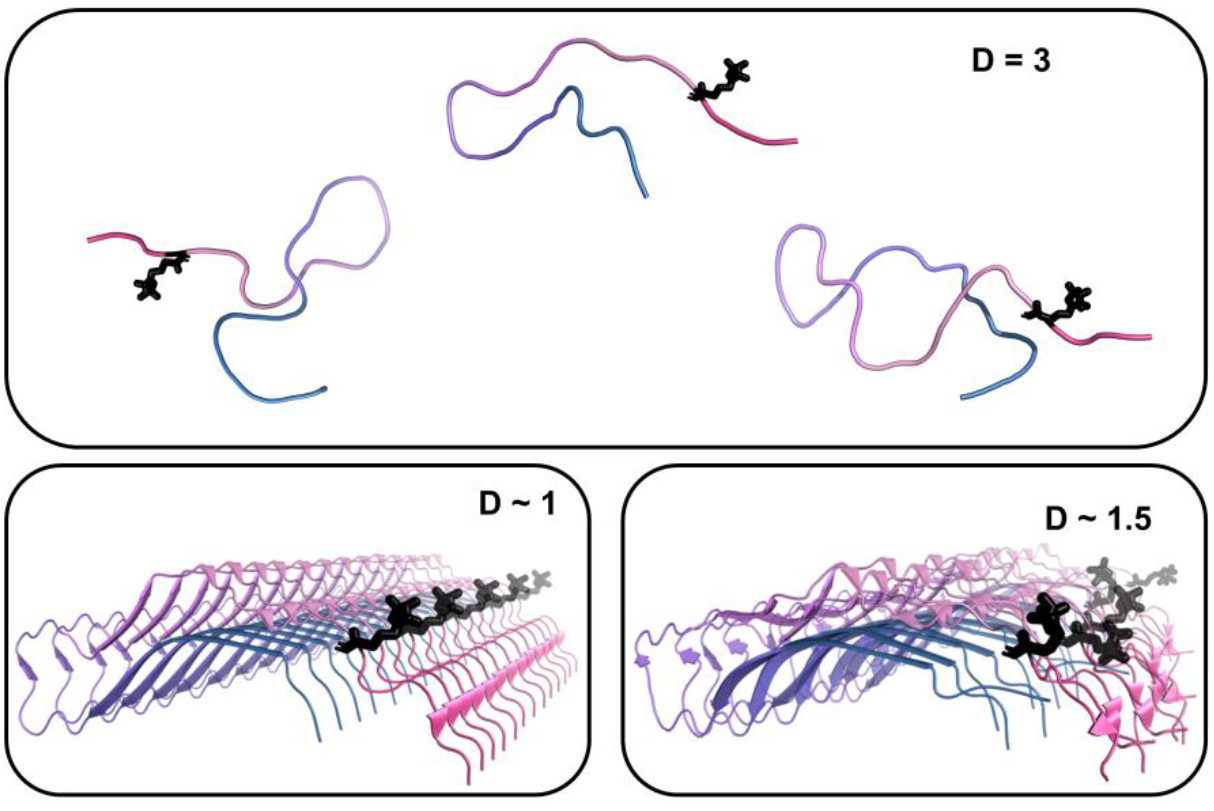

## 2 Introduction

Protein aggregation is involved in a large number of neurodegenerative disorders (NDs), including Alzheimer’s Disease (AD), Parkinson’s Disease and Huntington’s Disease. In these pathologies, proteins that are normally soluble or found in bound form to other biological constituents are found as insoluble fibrillar aggregates in different brain regions, also known as amyloid fibrils. They are highly ordered assemblies of proteins that stack in cross-β sheet structures (Fitzpatrick et al. 2013). These proteins are stabilized by a dense and highly organized network of hydrogen bonds and are therefore extremely stable. Strikingly, tau protein fibrils with subtle differences in the core fold and cross-β sheet arrangements are diagnostic hallmarks of different NDs, collectively referred to as tauopathies (Shi et al. 2021). However, the folding and assembly mechanism of tau proteins to recreate synthetic amyloid fibrils that adopt the disease fold and replicate prion-like competency is poorly understood (Broc et al. 2025). A pre-requisite for amyloid formation is the misfolding of proteins and the strict in register stacking of 2D folded protein segments along the fibril axis. Moreover, each monomer has to adopt precisely the same fold than the layer underneath in order to faithfully propagate a particular structure through nucleation/elongation. To effectively study the mechanism of amyloid formation, we need structural biology tools that can not only locate cross-β-sheet structures, but also yield insights into the packing and homogeneity of the amyloid fibril packing.

The main methods to investigate the structure of amyloid fibrils at high resolution are solid-sate NMR and cryoEM (Loquet et al. 2018; Scheres et al. 2023). NMR requires a large number of restraints as well as high homogeneity within the sample. It also heavily relies on intramolecular interactions, providing fewer details on how monomers are stacked along the fibril axis. CryoEM has provided enormous insights into tauopathy specific fibril structures in the last several years, mostly because of its capacity to resolve ex-situ fibrils (Scheres et al. 2023; Shi et al. 2021). However, CryoEM provides a view of only highly ordered and homogeneous fibrils and relies on helical reconstruction, where inter-layer packing must be identified and homogeneous. Ultimately, high-resolution structural biology methods provide limited information on how the proteins are packed along amyloid fibril axis.

Electron paramagnetic resonance (EPR) spectroscopy is a powerful method that can probe structure and dynamics of paramagnetic biomolecules and their assembly. For protein science (Jeschke 2012; Sahu et al. 2013), EPR has been used both in metalloproteins containing intrinsic paramagnetic metals and proteins labelled at select sites with paramagnetic probes. The pulse EPR method called double electron electron resonance (DEER) measures dipolar oscillation to extract the distribution of distances between pairs of electron spins, introduced by site-directed spin labeling onto target residues. DEER has been used to probe the intramolecular protein fold of tau within amyloid fibers, to show that *in vitro* tau fibril models differ from that of brain extracted tau fibers (Fichou et al. 2018b), or to reveal the selection of particular tau folding motifs during seeding (Fichou et al. 2018a; Meyer et al. 2016). DEER is most often used to obtain intramolecular distance distributions of 1.5 to 8 nm, thereby revealing protein conformation within the aggregate (Fichou et al. 2017; Schiemann et al. 2021).

When the paramagnetic centers are randomly distributed in the sample, the average of multiple oscillation frequencies leads to a feature-less decay of electron spin echoes (ESE). Milov and Tsvetkov have shown that the ESE decay function reflects both the density and spatial organization of the particles, i.e. whether they are in a straight line (1D) on a surface (2D) or in a volume (3D) (Milov et al. 2002; Milov & Tsvetkov 1997; Tsvetkov et al. 2008). One study reported that peptaibols integrated to a membrane surface exhibited a dipolar coupling from 2D like space, in contrast to the 3D signal obtained in solution (Milov et al. 2002). In another study, it was demonstrated that the local density of spin radicals randomly incorporated in a polymer could be derived from the ESE decay (Milov & Tsvetkov 1997). However, the ESE decay is not commonly used to characterize the structural properties of protein fibrils.

This study presents the ESE decay function as a promising approach to study protein aggregation that can inform on changes in the local density (i.e. aggregation) over time, and, concurrently, on changes in dimensionality when the protein transition from a soluble state (3D) to cross-beta sheet packing (2D assembly) polymerizing along a single dominant axis (1D). Specifically, we investigated how DEER can inform on the supramolecular arrangement of amyloid fibrils made of the intrinsically disordered protein, tau. We show that singly labelled tau proteins or peptides report on the local density in the assembly as well as their packing dimensionality. The approach provides a new method for quantifiying order and packing across self-assembled biological and synthetic supramolecular systems.

## 3 Results and discussions

### 3.1 Density and spatial arrangement of paramagnetic centers can be extracted from the DEER signal

Dipolar coupling between two spin pairs, typically measured by DEER, gives rise to an ESE oscillation that contains information on the distances between the spin pairs. These oscillations originate from coherence dephasing induced by dipolar coupling evolution. When oscillations are averaged over different frequencies the ESE evolution decays monotonously. In particular, in the case of randomly distributed and oriented paramagnetic centers (PC), one can show that the ESE decay takes the following form (Milov et al. 1998):

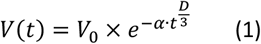

Where the parameter *D* reflects on the dimensionality of the space in which the PC evolves (*D* = 1 for a line, *D* = 2 for a plane and *D* = 3 for a 3D space) and *α* is a prefactor proportional to the local density of the PC and the pump efficiency. Note that Equation (1) is commonly used as the so-called background decay function of the DEER time domain data. Equation (1) is then divided from the acquired signal to remove contribution from inter-pair interactions, usually assumed to be randomly distributed.

In the 3D cases, e.g. for an ideal solution of PC, *α* takes the following expression (Salikhov et al. 1976):

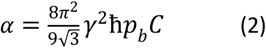

Where γ is the electron gyromagnetic ratio, ħ is the reduced Planck constant, *C* is the local concentration of PC and *p*_*b*_ is the probability of changing orientation of the spins excited by the pump pulse. i.e. the pump bandwidth.

When *D* is not equal to 3, *α* can be linked to the concentration using the following equation (Kutsovsky et al. 1990; Milov & Tsvetkov 1997), where the integral *I(D)* contains terms representing the dipolar modulation decays over time for a given dimension D and contribution from different spin orientations, and *k(D, C)* is a scaling function that contains the Euler’s function, *Γ*(*x*) to account for effective spatial dimension:

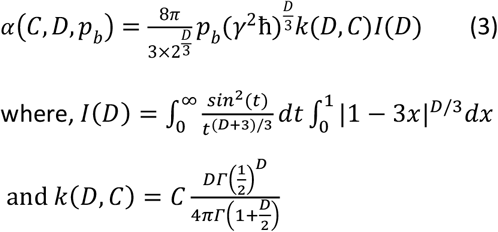

The notion of dimensionality might intuitively lead to the idea that the respective orientation of the spins with one another, along a line, within a plane or in 3D space, influence the ESE decay signal. However, under the conditions used in most DEER experiments at Q-band or lower field, the samples are amorphous and the powder averaging assumption is correct (i.e. no orientation selection). Rather, the space in which the PC evolves affects the distance distribution (independently of their orientation), which in turn determines the measured ESE signal. To demonstrate this, we calculated analytically the distance distribution of the nearest neighbor, *P*(*r*), of non-interacting particle randomly distributed in 1D, 2D and 3D spaces (Figure 1A) (Chandrasekhar 1943):

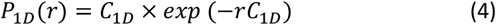

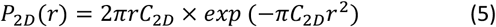

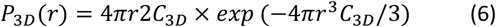

**Figure 1:**
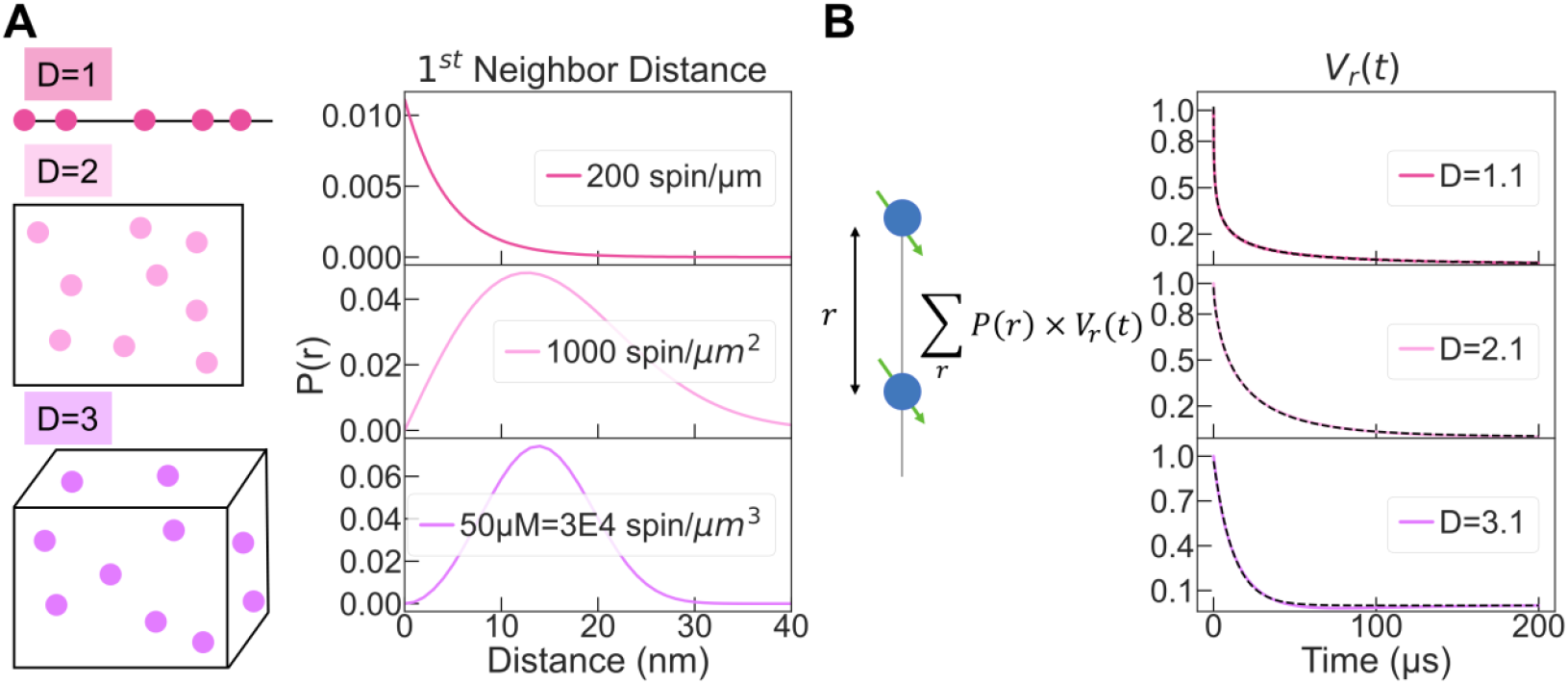
(A) First neighbor distance distribution can be analytically derived for PC randomly distributed in different space dimensions (line, plane and volume). (B) Computing the dipolar coupling for these distributions gives rise to a non-oscillating signal *V*(*t*) that can be fitted with Equation (1). The fitting output *D* parameter recapitulates the expected dimensionality.

Here, *C* is the spin concentration in the D-dimensional space.

Distributions (4)-(6) are plotted in Figure 1A for *C*_1*D*_, *C*_2*D*_ and *C*_3*D*_ equal to 212 μm^-1^, 1000 μm^-2^ and 60,221 μm^-3^ (corresponding to 10 μM), respectively. As expected, the shape of *P*(*r*) of the first neighbor is significantly different for the different dimensionalities. The DEER signal originating from these configurations were simulated as follow. The dipolar interactions calculated between two beads separated by a distance *r* were averaged over the distributions *P*_*nD*_(*r*) (Figure 1B). The obtained DEER signals are shown in Figure 1B and were fitted using Equation (1). The fit of the Dipolar dephasing signal from *P*_1*D*_(*r*), *P*_2*D*_(*r*) and *P*_3*D*_(*r*) lead to *D* of 1.1, 2.1 and 3.1. This result shows that the dimensionality *D* of the spin bath, as extracted from the fit of DEER decays, can be simulated by considering the first neighbor distribution *P*_*nD*_(*r*). Following the same procedure, different spin concentrations were simulated in each dimensionality (Figure S1). As predicted by Equation (1)-(3), *D* is independent of the concentration and solely depends on the shape of the distribution while *α* varies with the spin concentration. Overall, these simulations demonstrate that the DEER decays contain information about the spatial arrangement and the density of spins in the system, and that this information can be extracted from the time-domain DEER data.

We experimentally tested the approach by measuring DEER of the free radical 4-OH-TEMPO at different concentrations (Figure 2). This sample represents the ideal case of uniform spin bath evolving in a volume. The signals for each concentration were fitted with Equation (1) with *V*_0_, *α*, and *D* as free parameters. The fitting parameters are shown in Figures 2B,C. As expected, we see that *D* is close to 3 for all concentrations but 1 mM. At this higher spin concentration, it is likely that the excluded volume of 4-OH-TEMPO can no longer be neglected and the assumption of a random distribution does not hold (Kattnig et al. 2013). The *α* parameter is found to be exactly proportional to the concentration of PC (Figure 2C). From Equation (2), it follows that the efficiency of the pump pulse *p*_*b*_ is concentration independent under our experimental condition, and is found to be 0.73.

**Figure 2:**
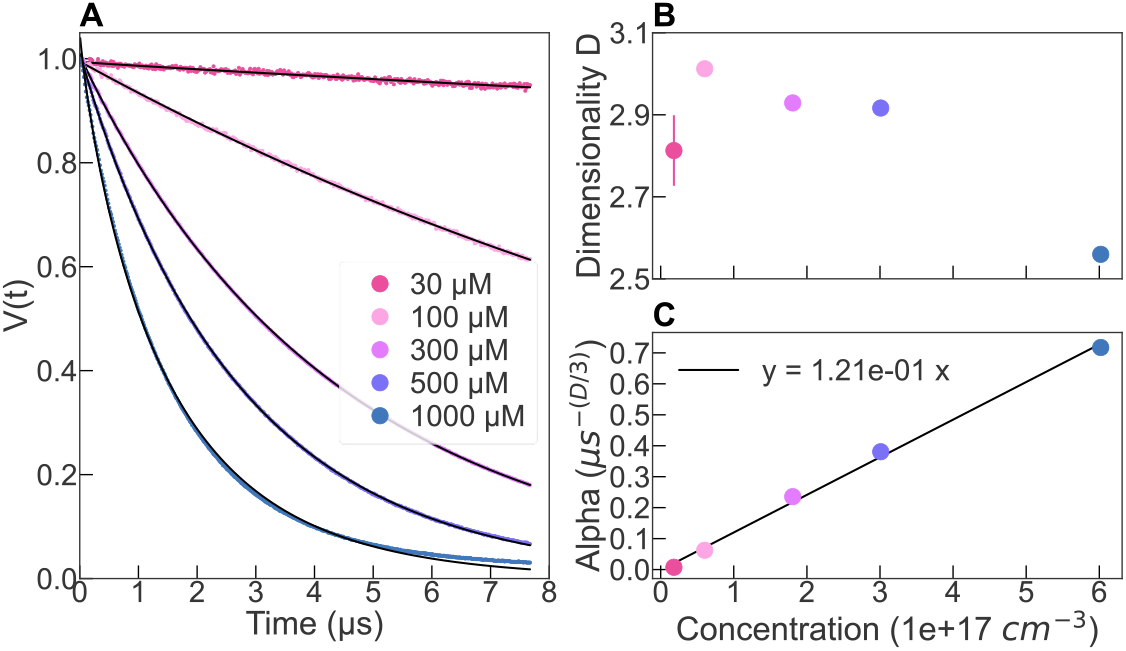
(A) ESE decay of 4-OH-TEMPO in solution fitted with Equation 1 (black solid line). (B) Output parameters of the fit. Concentration is in number of particles per cm^3^ (1 µM = 6e14 cm^-3^).

One direct application of this relation is the possibility to precisely determine the local spin concentration of a sample by measuring the ESE decay. Similarly, when making typical measurements of intra molecular distances of a doubly labelled molecule, one can evaluate the concentration of molecules by looking at the slope of the background, which can be fit at times larger than the last oscillation of the intra molecular DEER signal.

Furthermore, when using a free radical solution of a known concentration, one can use Equation (2) to optimize the efficiency of the pump pulse. For instance, in Figure S2, we have increased the power output of the pump pulse from 20 % to 100%, while keeping a fixed bandwidth of 40 MHz. Figure S2 shows that above 60% of pump power, the value *α*, and therefore *p*_*b*_, starts to plateau, implying that increasing the power above 60% in the same experimental configuration should not improve the signal. Rather, one can for instance increase the bandwidth of the pump pulse, while using the maximum power, so that a larger part of the spectrum can be excited, thereby providing a better signal to noise ratio (Figure S2). For instance, we have increased the pump spectral bandwidth from 40MHz to 130MHz, while using 100% of the pump pulse output (Figure S3). We see that *α* increases as the bandwidth increases, showing that the pump efficiency is higher when increasing the bandwidth.

### 3.2 DEER of singly labelled protein reveals protein condensation and alignment in the amyloid core

We prepared amyloid fibrils of tau by incubating the protein with the polyanion heparin. We used a C-terminus truncated 0N4R tau spanning residues 244-441, a 187-residue-long protein referred to as tau187, that forms amyloid fibrils upon addition of heparin (Pavlova et al. 2016). Fibril formation was confirmed by Thioflavin T (ThT) fluorescence (Figure 3D) and negative stain transmission electron microscopy (TEM) (Figure 3C). Amyloid fibrils are linearly stacked assemblies of proteins folded in approximately 2D folds forming cross-β sheets along the fibril axis. Each layer of the tau protein in the fibril is spaced by 4.7 Å (Figure 3B). While this 4.7 Å gauge between protein layers is constant for all amyloid fibrils, the arrangement within a given layer of the protein varies greatly depending on the protein and the aggregation conditions (Figure 3B).

**Figure 3:**
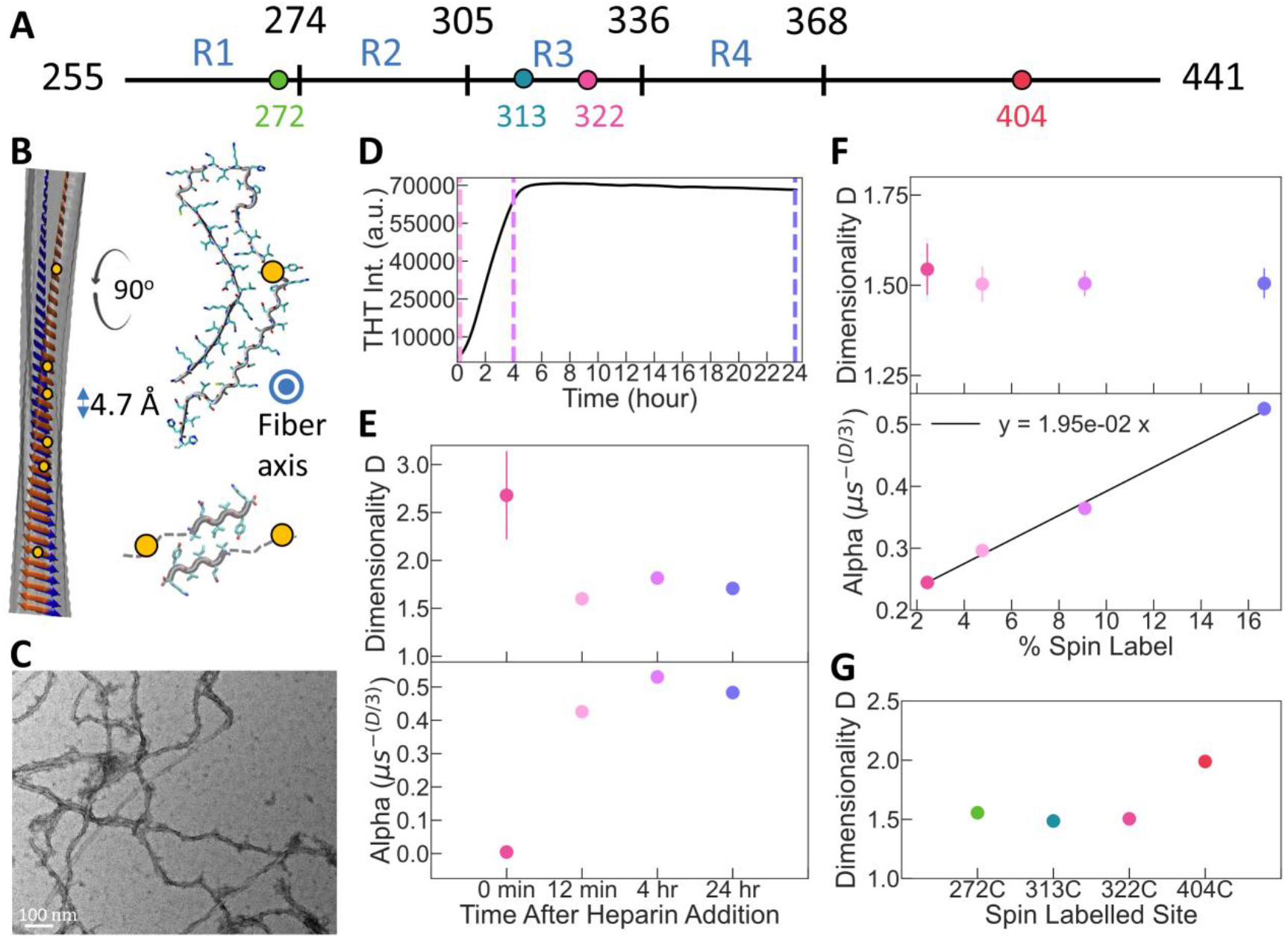
DEER signal probe protein condensation and alignment during amyloid formation. (A) A fragment of the tau protein, referred to as tau187, used to form amyloid aggregates. (B) Amyloids aggregates are elongated filaments in which each protein monomer is stacked with a cross-beta sheet structure. Within each layer, a protein can take different conformations: model of full-length tau fibers made with heparin (derived from PDB 6qjh; upper structure) and of tau peptide VQIVYQ (derived from PDB 2ON9; lower structure). (C) TEM of tau187 aggregates. (D) THT fluorescence intensity of tau187 aggregation. The dash lines represent the times at which samples were flash-frozen for analysis. (E) *D* and *α* parameters extracted from fit at each time points. (F) *D* and *α* parameters for filaments made with different SL-tau:noSL-tau ratios. The x-axis represents the percentage of total protein that is labelled. SL-tau protein of labelled to unlabeled tau187. (G) *D* and *α* parameters for filaments made with tau187 labelled at different positions, for a SL-tau:noSL-tau ratio of 1:10. The raw DEER signals and their fits are presented in Figure S4,S5 and S6.

We incubated a mixture of labelled (SL-) and unlabeled (noSL-) tau proteins with heparin that results in the complete formation of tau fibrils within about 6 hours (Figure 3D). The variation of the ratio of SL-tau and noSL-tau allows effective variation of the spin density along the amyloid fibers. First, we measured DEER of tau fibrils made with ratios of SL-tau:noSL-tau of 1:5, 1:10, 1:20 and 1:40 (Figure S4). The output parameters of the fit are shown in Figure 3F. The parameter *α* is found to be linear with the ratio SL-tau:noSL-tau, i.e. with increasing spin label concentration along the fiber axis (Figure 3F bottom). This linear dependence of *α* is in good agreement with Equation (3) and the understanding that the pump bandwidth *p*_*b*_ is independent of the spin concentration (Figure 2C). In other words, *α* is a direct measure of the concentration of spin labeled protein according to Equation (3), knowing *p*_*b*_ from the reference experiment (here 0.73). The dimensionality *D* is consistently found to be 1.51 +/−0.02, independent of the spin concentration (Figure 3F top). This reduced dimensionality reflects the alignment of the labelled residue along the fibril axis. Its quantitative interpretation is further investigated in Section 3.4.

### 3.3 Probing amyloid core location and aggregation kinetic

Next, we repeated the experiment with a ratio SL-tau:noSL-tau of 1:10, using tau labeled at different sites along the protein, specifically residues 272, 313, 322 and 404 (see Figure 3A). Sites 272, 313 and 322 are expected to be part of the amyloid core, while 404 is part of the so-called fuzzy coat, the part of tau that remains disordered even in the fibers. Remarkably, all sites in the amyloid cored exhibited *D*=1.55 +/−0.03 while site 404 in the fuzzy coat shows *D*=2 (Figure 3G). The fuzzy coat is expected to dynamically coat the fibril surface, and hence the spin label is expected to be distributed on the exterior fibril surface, not in solution, rendering the spatial distribution of spin labels on the fuzzy coat as 2D. It demonstrates that the dimensionality can differentiate between the amyloid-forming region and the fuzzy coat, and hence is a sensitive tool to map the amyloid-core regions of protein aggregates.

Finally, we recorded the DEER signal before (monomeric tau), 12min, 4h and 24h after heparin addition (Figure S6). The kinetic of the aggregation reaction under these conditions was followed by ThT fluorescence (Figure 3D), which reports on the quantity of cross-β structures and hence on the fibril quantity in a sample. As shown by Figure 3D, the fluorescence intensity has barely increased at 12 min, while it plateaus to its maximum value around 6h. Figure 3E shows the extracted parameter *D* and *α* from the fit of the data acquired at different times. We see a clear increase in *α* from 0.004 before heparin addition to 0.4 at 12 min after heparin addition, revealing a drastic increase of the local spin density. Only slight increase in *α* is seen between 12 min and 4h, indicating that most of the protein condensation takes place in the first 12 minutes. *D* on the other hand decreases from ~3 before heparin addition to 1.6 within 12 min after heparin addition, and then stays around this value at longer times. Note that the error bar is significantly higher for tau monomeric because the *α* is small (i.e. low concentration), which leads to more ambiguous fitting but still around the expected *D*=3. These measurements show that within minutes after heparin addition, i.e. before forming cross-β network leading to measurable ThT signal (Figure 3D), the tau proteins agglomerate together (large *α*) in a conformation that might be close to 1D stacked amyloid structural assembly already (*D* close to 1.5). This is in good agreement with previous finding that tau and heparin immediately form metastable oligomers that convert to amyloid fibrils (Fichou et al. 2019). These results explicitly demonstrate that the early, ThT-active, oligomers have already adopted 1D stacked amyloid arrangements.

### 3.4 Dimensionality can be linked to fibril geometry and packing

To help link the derived parameters from the ESR of DEER (*α,D*) with protein fibril geometry, we modeled the amyloid fibril with a line of beads, spaced by 0.47 nm, each of which has a probability *p* to be considered as a paramagnetic bead (Figure 4A). *p* corresponds exactly to the ratio of labelled:unlabeled protein used to prepare the fibrils. From this line of beads, we computed the distance distribution *P*(*r*) of the first neighbor of PC. Note that this numerical simulation gave strictly the same distribution that the analytical distribution given in Equation (4) for *C*_1*D*_ = *p*/4.7, as shown in Figure S7. The *P*(*r*) was then used to simulate the corresponding ESE decay, which was fitted with Equation (1). In this linear configuration, the fit gave *D*=1 and *α* depends linearly on *p* (Figure 4B), as found experimentally in Figure 3F. We used this modeling framework to simulate different configurations described in the next paragraph.

**Figure 4:**
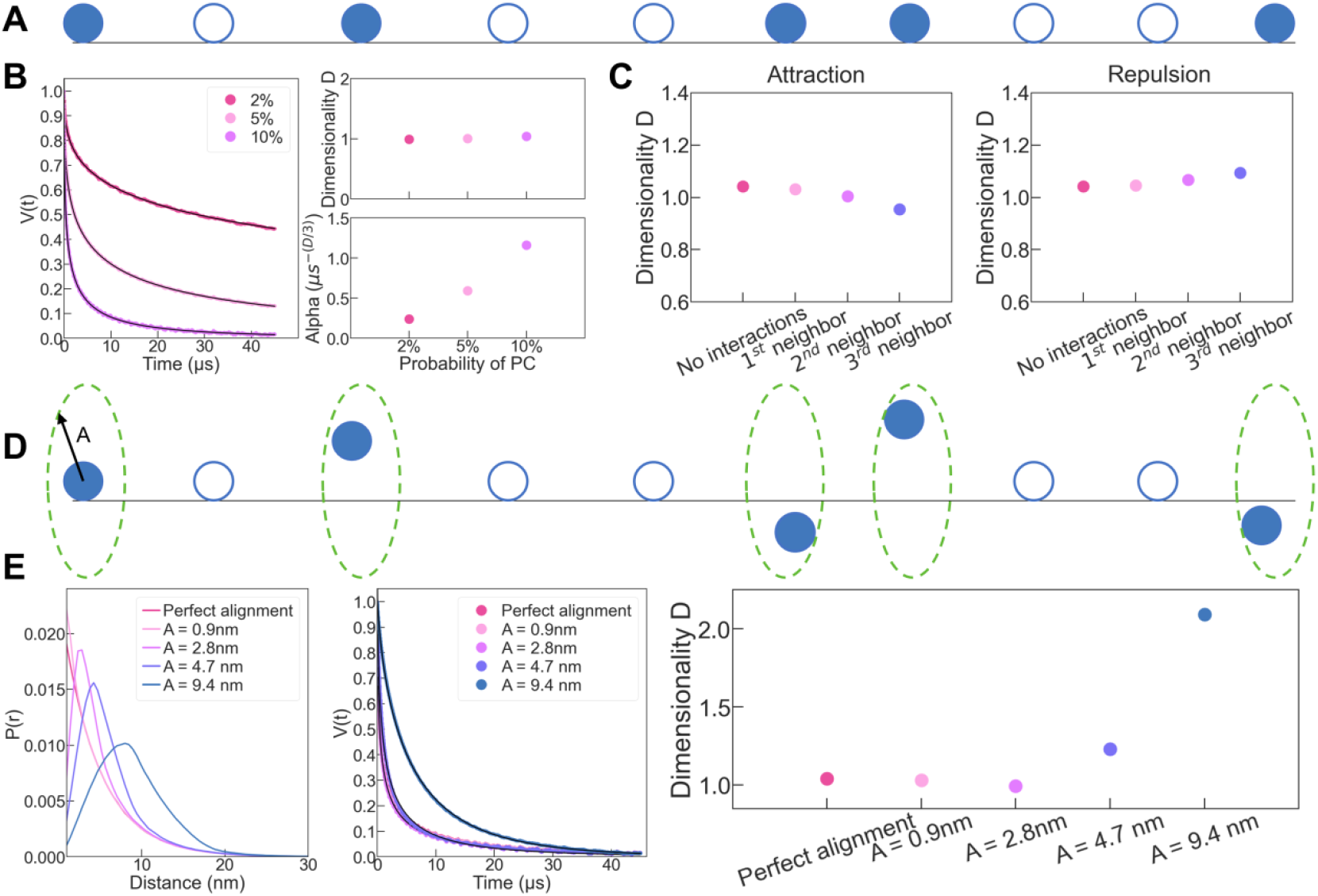
(A) amyloid filaments were modelled with a line where PC (blue balls) has a probability *p* to exist at each discrete locations spaced by 4.7 Å (blue circle). First neighbor distance distribution of PC was numerically calculated from this model and used to compute dipolar signal (B) for *p*=0.02 (dark pink) *p*=0.05 (light pink) and *p*=0.1 (lilac). *D* and *α* parameters were extract from fitting with Equation (1). (C) Fitting output parameter *D* when *p* is twice more (attraction) or less (repulsion) important in the vicinity of a PC. The vicinity extends to the 1^st^ (light pink), 1^st^ and 2^nd^ (lilac) or 1^st^, 2^nd^ and 3^rd^ (blue) neighbors. (D) Misalignment is introduced by allowing each PC to move within a disk of radius, *A*. (E) The resulting distance distributions are converted into dipolar evolution times that are fitted with Equation (1). (C) and (E) are generated with *p*=0.1.

Tau amyloid fibrils are composed of *in register* β*-*sheet (Margittai & Langen 2004; Zhang et al. 2019), meaning that each residue is packed on top of each other along the fibril (Figure 3B). Consequently, each spin label should be aligned in a linear configuration along the fibril over 100s of nm (Figure 3B). This being an ideal case, we consider three features that could create a deviation from this situation. (i) The sample is composed of both monomers and amyloid fibrils. (ii) the spins are not randomly distributed along the fibril axis. (iii) The local flexibility of the label and/or defects in the packing leads to a misalignment. We ruled out (i) experimentally by showing that ultracentrifugation of the fibril samples at 100,000g revealed that less 5% of the protein remains soluble, quantified by UV-VIS of the supernatant. In addition, mathematical modelling shows that the dimensionality parameter has little sensitivity to the presence of soluble particles (Figure S8). We investigated next how the dimensionality would reflect on the features (ii) and (iii), with the specific goal to interpret the value *D* ~ 1.5 found experimentally (Figure 3).

If spin labelled proteins interact more (attraction) or less (repulsion) preferentially with themselves compared to with unlabeled protein, the spin would not be randomly distributed along the fibril axis (feature (ii)). While attraction is less likely, repulsion is possible through steric hindrance of the spin label (MTSL, 260 Da). Using our model, we implemented repulsion and attraction over 1 to 3 neighbors, by doubling (attraction) or dividing by 2 (repulsion) the probability *p* at position n+1 to n+3, if a spin is present at position n. As shown in Figure 4C, we found little effect on the dimensionality, suggesting that interaction between spin labels is not the basis for the experimentally measured *D*~1.5.

Although tau fibrils are overall enriched in β sheets structures, it is unclear if they present some local defect that creates protein flexibility and structural heterogeneity (feature (iii)). This might be amplified by the presence of a relatively large spin label (260 Da). Therefore, the protein flexibility together with the spin label rotation would create a deviation from a perfect linear alignment. We simulated this effect by allowing each spin on the line to be randomly located inside a disk of radius *A* perpendicular to the line (Figure 4D). Figure 4E shows the computed distance distributions and the corresponding dipolar evolution time, which is fitted with Equation (1) for different amplitude *A*. A dimensionality of 1.5 is reached when the fluctuation is between 5 and 10 nm around the ideal line. This finding reveals that the dimensionality of 1.5 consistently found experimentally for tau filament likely reveal a misalignment.

### 3.5 Revealing tau amyloid fibril mispacking that can be mitigated with templating

The modeling of the previous section pointed to misalignment of the labelled residues as a potential origin of the experimental dimensionality of 1.5. To verify this experimentally, we used a short tau fragment of 16 amino acids (residues 295 to 313 of tau full length; sequence in Figure 5A) that we refer to as tau16. This peptide is highly similar and only 3 residues shorter than another 19-residue tau fragment, R2R3-P301L, that has already shown to form neat amyloid fibrils of sufficiently high quality to yield high-resolution cryoEM structure (Vigers et al. 2025). By further reducing the length of the protein, we expect to reduce the chance of mispacking and deviation from the canonical cross-beta sheet structure, as previously shown for 6-10 residue long tau peptides (Sawaya et al. 2007; Seidler et al. 2018). We labelled this tau16 peptide at position V313C and prepared a mixture of 1:10 SL-tau16:noSL-tau16. Similar to tau187, we incubated tau16 with heparin in order to form amyloid filaments (Figure 5A). The ESE decay of tau16 filaments was recorded and fitted with Equation (1), leading to *D*=0.8 (Figure 5D). The finding that *D* is closer to 1 as compared to longer version of tau strongly suggest that *D*=1.5 found for tau187 reveal an imperfect packing of the in-register cross-beta sheet structures.

**Figure 5:**
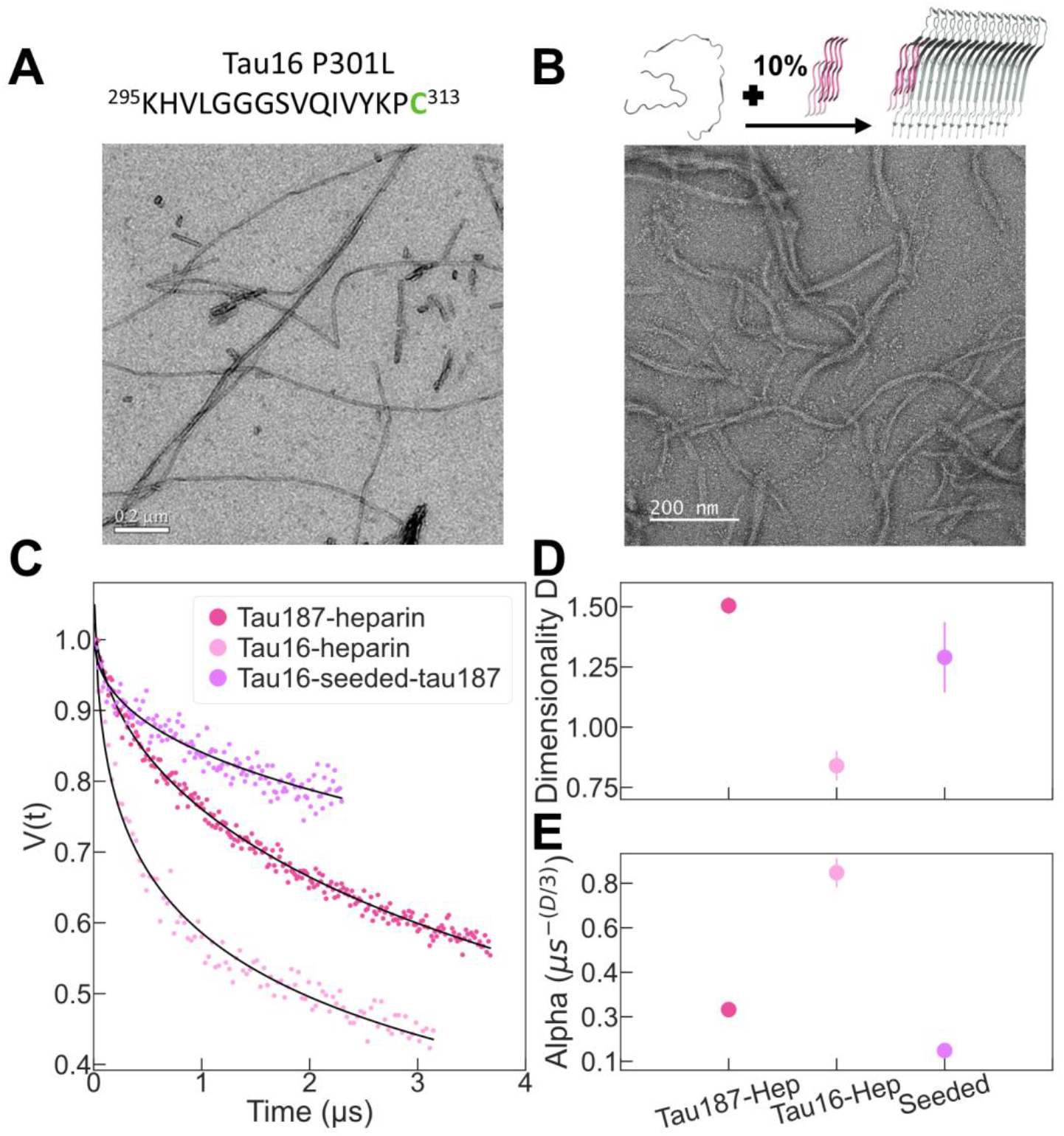
We compare heparin-induced tau187, heparin-induced tau-16, and tau16 seeded tau187. (A) tau16 peptide can aggregate into amyloid filaments expected to be well packed. (B) Tau16 filaments can be used to seed tau187 peptide into amyloid filaments. (C) The DEER signal of tau filaments are fitted with Equation (1) (black line) to extract (D) the dimensionality and the (E) *α* parameters.

Heparin-induced filaments was already proposed to be mispacked (Fichou et al. 2018b), which is compatible with the imperfect packing leading to *D*=1.5 found for tau187. Because tau is known to exhibit prion-like properties (Duan et al. 2024), we hypothesized that having a well ordered template would limit such misalignment. To test this experimentally, we used tau16 filaments that show perfect packing as the “seed” to template tau187 aggregation, instead of using heparin as the cofactor. We made the tau16 seeds by incubating tau16 monomers with heparin followed by sonication to break the filaments. We then seeded tau187 (1:10 SL-tau187:noSL-tau187 mixture, with spin label of SL-tau187 at site C322) with tau16 seeds at 10% mass of the total tau protein material in the mixture. The efficiency of seeded aggregation of tau187 was monitored by ThT and TEM (Figure 5B and S9). The DEER signal of tau16-seeded-tau187 filaments was recorded and fitted with Equation (1), giving *D*= 1.29 +/−0.15 (Figure 5D). The finding that *D* = 1.3 for tau16-seeded-tau187, and hence closer to 1 than *D* = 1.5 found for heparin-induced tau187 filament suggests that there is a templating mechanism where well-packed seeds can improve the packing quality of the in-register cross-beta sheet structures of heparin-induced tau187.

### 3.6 Probing protofilament composition

Amyloid filaments can be composed of one or several protofilaments (Figure 3B). The protofilament arrangement of tau amyloids is one of the key identifying structural properties, besides the protein fold, that differentiate between different tau pathologies (Shi et al. 2021). Our aim was to test whether the proposed approach is sensitive to the protofilament composition of tau filaments. Because each protofilament is composed of one protein per layer of 4.7 Å thickness, the number of protofilament should correlate with the spin density per length of filament, which is measured by the *α* parameter, as shown in Figure 3F. We compared the ESE decay of tau16 and tau187 filament made with heparin with the same ratio SL-tau:noSL-tau of 1:10 (Figure 5).

We found that *α* is more than 2 times higher for tau16 (*α* = 0.8) than for tau187 (*α* = 0.3) (Figure 5E), demonstrating much greater spin density per fibril layer in the former case. The result suggests that tau16 filaments are composed of at least two monomers per fibril layer and tau187 filaments of one monomer per layer. This is consistent with the cryoEM structure of tau-heparin filaments that showed mostly single-protofilament amyloids ((Zhang et al. 2019) and Figure 3B) while shorter peptides needs typically 2-4 monomers per layer in order to form a steric zipper ((Sawaya et al. 2007; Vigers et al. 2025) and Figure 3B). Note that the same conclusion is reached if *D* is constrained to 1 for both datasets (Figure S10). Thus, we show here that our approach can assess the number of protofilament present in amyloid filaments.

## 4 Conclusions

We have developed a novel framework to probe spatial arrangement of protein aggregates with DEER experiments. We showed that the dipolar interaction from singly labelled proteins can be used to gain information on supramolecular packing of amyloid fibrils, which are disease-relevant protein assemblies. We specifically focused on the tau protein that can take disease-specific folds in a class of disorders called tauopathies. The linear-like configuration of amyloid filaments gives rise to a reduced effective dimensionality, which can be used to locate the amyloid core region and follow the formation of intermediates during aggregation. Furthermore, a quantitative analysis of the dimensionality showed that this parameter can reveal the packing ordering of amyloids fibrils. Specifically, we show that heparin fibrils made of long versions of tau are mispacked, in contrast to shorter fragments that form seeding-active high-quality fibrils exhibiting lower conformational variability. Seeded aggregation was further shown to achieve superior templating and packing of the tau proteins to fibrils, as manifested in a reduced dimensionality. Finally, we showed that the local spin density obtained through the *α* parameter can be used to assess the number of protofilament composing the filaments. Overall, we have proposed and benchmarked a new method to characterize protein supramolecular organization of amyloid filaments, which is a critical structural characteristic of amyloid fibrils. The method can be applied to any amyloid-forming protein functionalized with a single spin label and provides unique information complementary to conventional methods such as NMR and cryoEM data.

## 5 Material and methods

### Protein expression, purification and aggregation

An N-terminal truncated form of 2N4R tau, referred to as tau187, containing the microtubule binding domain (residues 255-441 from the longest human tau isoform 2N4R with a 6× His-tag at the N-terminus) was used for *in vitro* studies. Tau187 contains two native cysteines, at residues 291 and 322. Several mutants of tau187 were prepared in order to control the position of a single cysteine, used for spin labelling. These mutants are prepared by site-directed mutagenesis and are referred to with the position of the cysteine: tau187-322C contains C291S mutation, tau187-cysless contains C291S and C322S, tau187-272C contains C291S, C322S and G272C, tau187-313C contains C291S, C322S and V313C and tau187-404C contains C291S, C322S and S404C. The expression and purification of tau187 have been previously reported (Fichou et al. 2018b,a; Pavlova et al. 2016) and is recalled here. E. coli BL21 (DE3) cells were transfected with constructed DNA variants and stored as frozen glycerol stock at −80 °C. Cells from glycerol stock were grown in 10 mL luria broth (LB, Sigma Aldrich, L3022) overnight and then used to inoculate 1 L of fresh LB. Growth of cells were performed at 37 °C, 200 rpm with addition of 10 μg/mL kanamycin (Fisher Scientific, BP906) until optical density at λ = 600 nm reached 0.6–0.8. Expression was induced by incubation with 1 mM isopropyl-ß-D-thiogalactoside (Fisher Bioreagents, BP175510) for 2–3 h. Cells were harvested with centrifugation at 4500 g for 20 min. Harvested cells were resuspended in lysis buffer (Tris-HCl, pH = 7.4, 100 mM NaCl, 0.5 mM DTT, 0.1 mM EDTA) added with 1 Pierce protease inhibitor tablet (Thermo Scientific, A32965), 1 mM PMSF, 2 mg/mL lysozyme, 20 μg/mL DNase (Sigma, DN25) and 10 mM MgCl_2_ (10 mM), and incubated on ice for 30 min. Samples were then frozen and thawed for 3 times using liquid nitrogen, then centrifuged at 10,000 rpm for 10 min to remove cell debris. 1 mM PMSF was added again and samples were heated at 65 °C for 12 min and cooled on ice for 20 min. Cooled samples were then centrifuged at 10,000 rpm for 10 min to remove the precipitant. The resulting supernatant was incubated overnight with Ni-NTA resins pre-equilibrated in buffer A (20 mM sodium phosphate, pH = 7.0, 500 mM NaCl, 10 mM imidazole, 100 μM EDTA). The resins were loaded to a column and washed with 20 mL of buffer A, 25 mL buffer B (20 mM sodium phosphate, pH = 7.0, 1 M NaCl, 20 mM imidazole, 0.5 mM DTT, 100 μM EDTA). Purified protein was eluted with 15 mL of buffer C (20 mM sodium phosphate, pH = 7.0, 0.5 mM DTT, 100 mM NaCl, 300 mM imidazole). Eluents were analyzed by SDS-PAGE to collect the pure fractions. Proteins were then buffer exchanged into DTT-free working buffer (20 mM HEPES, pH 7.0). Tau16 peptides (KHVLGGGSVQIVYKPC and KHVLGGGSVQIVYKPV) was synthetized (Genscript, custom peptide synthesis).

### Protein labelling and radical preparation

Powder of 4-OH-TEMPO (Sigma-Aldrich; 176141) was dissolved in DI water to reach 10mM solution. From this stock, lower concentration solutions were prepared by dilution. Tau187 was spin-labelled using MTSL ((1-Acetoxy-2,2,5,5-tetramethyl-δ-3-pyrroline-3-methyl) Methanethiosulfonate) purchased from Toronto Research Chemicals. Prior to labelling, samples were treated with 5 mM TCEP, which was removed using a PD-10 desalting column. Then, 10× molar excess MTSL to free cysteine was incubated with the protein at room temperature overnight. Excess MTSL was removed using a PD-10 desalting column. After labelling, the buffer was exchanged into D2O-based buffer (20mM HEPES pH 7.0, 100mM NaCl) for subsequent EPR experiments. The same procedure was used to labelled tau16, except that PD minitrap G-10 columns (cytivia) were used instead of PD-10 columns. Labelling efficiency, defined as the molar ratio of tethered spin labels over the cysteines, was measured to be between 60% and 65% for all mutants.

### Protein aggregation

The labelled cysteine-containing tau187 mutants were mixed with tau187-cysless mutant (i.e. containing C291S and C322S mutations) with a molar ratio tau187-labelled:tau187-cysless of 1:10, except when otherwise stated. All tau16 peptide was ordered from GenScript. The labelled cysteine-containing tau16 peptide (KHVLGGGSVQIVYKPC) was mixed with tau16 fragment mutant (KHVLGGGSVQIVYKP) with a molar ratio tau16-labelled:tau16 of 1:10. The total protein (peptide) concentration was typically 300 μM. Heparin (Galen laboratory supplies, HEP001) was then added to a tau187(tau16):heparin molar ratio of 4:1. The solution was then incubated for at least 24h at 37C, before DEER sample preparation.

The tau16 seeds were prepared using 260 μM tau16 fragment (KHVLGGGSVQIVYKP), 65 μM heparin, and 100 mM NaCl in 20mM HEPES, pH 7 buffer, then incubated for 40 hours at 37C. The seeds were fragmented using a cup-horn sonicator with 80% power for 2 minutes. The labelled cysteine-containing tau187 mutants were mixed with tau187-cysless mutant with a molar ratio tau187-labelled:tau187-cysless of 1:10. Then 10% mass of sonicated tau16 seeds were added to the tau187 peptide mixture and incubated for 24 hours. The seeding reaction was done with total peptide concentration of 50 μM and 1000 μL total volume in 20 mM HEPES, pH 7, 100 mM NaCl buffer.

### TEM

10 μL of was deposited on a copper grid (FCF-300-Cu) cleaned with plasma for 20 s. Samples were left for 1min, then blotted then rinsed with water for 1min. Finally, 10 μL 1.5 w/v % uranyl acetate was added. Grids were then imaged with a JEOL JEM-1230 (JEOL USA, Inc).

### Double Electron Electron Resonance (DEER)

28 μL of the sample solution were mixed with 12 μL D_8_-glycerol before transferring to a quartz tube (2 mm i.d.) and freezing using liquid nitrogen.

Four-pulse DEER experiments were carried out at 50 K using the Q-band Bruker E580 Elexsys pulse EPR spectrometer operating at ~34 GHz and equipped with a 300 W TWT amplifier. The following DEER pulse sequence was used: π_obs_/2 – τ_1_ – π_obs_ – (t-π_pump_) – (τ_2_-t) – π_obs_ – τ_2_ – echo. Observe pulses were rectangular with lengths set to π_obs_/2 = 10-12 ns and π_obs_ = 20-24 ns. A chirp pump pulse was used with a length of 20-24 ns and a frequency width of 60 MHz. The observe frequency was 90 MHz higher than the center of the pump frequency range. τ_1_ was set to 180 ns. The last 200 ns of the DEER data were cut before analysis. DEER data were accumulated for 12-24h.

### Fitting procedure

Fitting of the DEER signal with Equation (1) was performed with the lmfit package for python (lmfit.github.io/lmfit-py/), using a Levenberg-Marquardt minimizer. Error bar reported for *α* and *D* parameters represent are extracted from the output covariance matrix.

### Thioflavin T (ThT) fluorescent assays

Samples were mixed with 20 μM ThT and 20 μl of this mixture were placed in a 384-well microplate (Corning 3844). ThT fluorescence was measured in a plate reader (Biotek Synergy 2) at 37 °C without shaking.

### Simulation of distance distribution

A 1D matrix M was filled by 0 or 1 with a probability of (1-*p*) and *p*, respectively. The Matrix contained N=5E7 elements or more. For every 1 found in the matrix, the distance D to the nearest 1 (i.e the difference in their matrix indexes) was stored in a list L. The list elements were multiplied by 4.7 and gathered into a histogram of 400 bins to represent the simulated *P*(*r*) in Angstrom. The histogram was normalized so that its area is 1. Figure S7 verifies that this simulation procedure matches the analytical Equation (4) of the manuscript. **Simulating label interaction**: Each time an element of M was filled with 1, the nearest elements (over 1,2 or 3 elements) were filled with a penalty Pe so that the probability to place a 1 in these elements was *p*+Pe. Pe was positive to simulate label attraction and negative to simulate label repulsion. **Simulating imperfect alignment:** the distance D between nearest 1 in the matrix was considered to represent the distance between two disks of radius F. The labels were considered to take a random position on these disks. The effective distance *d* stored in the list L used to build the *P*(*r*) was therefore:

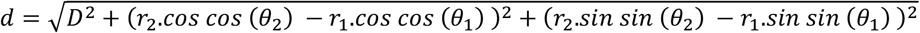

Where θ_*1*_, θ_*2*_ are random numbers in [0,2π[ and *r*_*1*_, *r*_*2*_ are random numbers in [0,F[.

### Simulation of dipolar signal from distance distribution

Simulations of dipolar dephasing between two electron spins were performed using Spin Evolution ((Veshtort & Griffin 2006) at Q-band frequency (35 GHz). The spin system consisted of two electron spins (pump and probe channels), and the coupling interaction was modeled using only the secular part of the dipolar coupling. Non-secular terms were neglected based on the large g-anisotropy at Q-band, which truncates such interactions effectively. The inter-spin distance distribution was incorporated using Cartesian coordinates from an external file, and dipolar couplings were calculated accordingly. The system employed no relaxation. The initial density matrix was set as coherence of an electron spin, and the observable was time evolution of the same coherence, in order to capture its evolution to dipolar interaction with the probe spin at 5 ns intervals. To account for spatial orientation of dipolar tensor in an external magnetic field, powder averaging was performed using an angular grid.

## Acknowledgement

YF thanks the European Research Council (Grant 101040138), the Fondation Vaincre Alzheimer and the Federation of European Biochemical societies for their financial support. KT and SH thank the Rainwater Charitable Foundation (RCF) for support through the Tau Consortium (Grant SB190039) and the National Institute of Health (Grant R01AG056058) for their financial support.

## Data availability

The datasets are organized by figure number and available at **10.5281/zenodo.18460124.**

