## Supplemental Information for "DEER of Singly Labelled Proteins to Evaluate Supramolecular Packing of Amyloid Fibrils"

### Supplementary figures

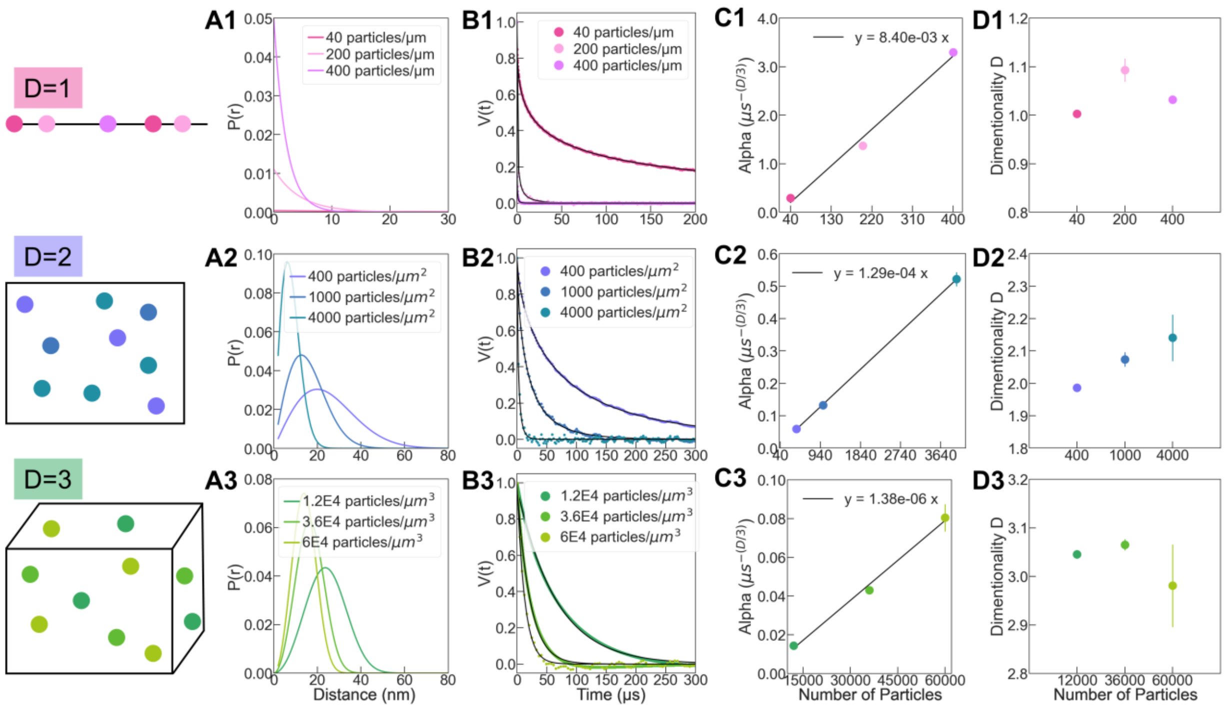


Figure S1: Distribution of particle first neighbor distances (A) computed from equation (4)-(6) were used to simulate the corresponding dipolar signal (B). Fitting of the latter signal with Equation (1) shows that the $D$ parameter (D) depends solely on the dimension of the particle bath while the alpha parameter (C) depends solely on the concentration of these particles.

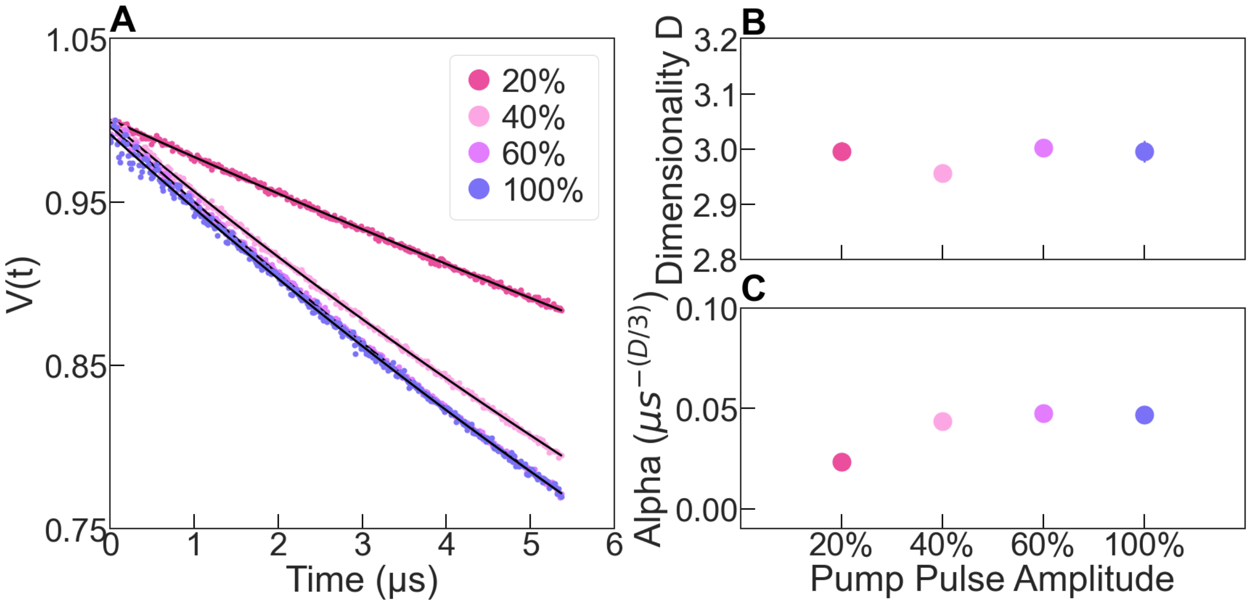

Figure S2: DEER experiment was performed on a solution of 100 µM OH-TEMPO. The power of the pump pulse was varied from 20 % to 100 %, at constant bandwidth of 40 MHz. The alpha parameter, which reflects the efficiency of the pump pulse (parameter $p_b$ in Equation (2)) show that 60% power was sufficient to reach optimal pumping.

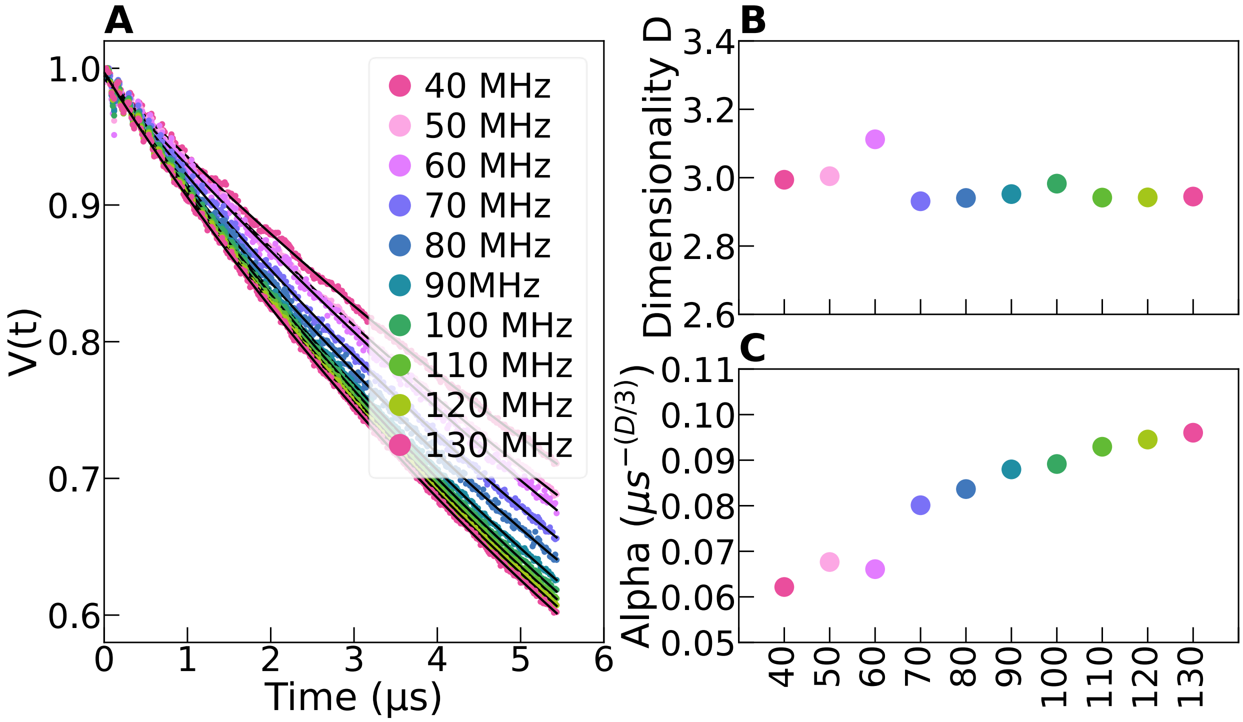


Figure S3. DEER experiment was performed on a solution of 100 µM OH-TEMPO. The bandwidth of the pump pulse varied from 40 MHz to 130 MHz. The alpha parameter, which reflects the efficiency of the pump pulse (parameter $p_b$ in Equation (2)) shows that increasing pump bandwidth increase pump efficiency.

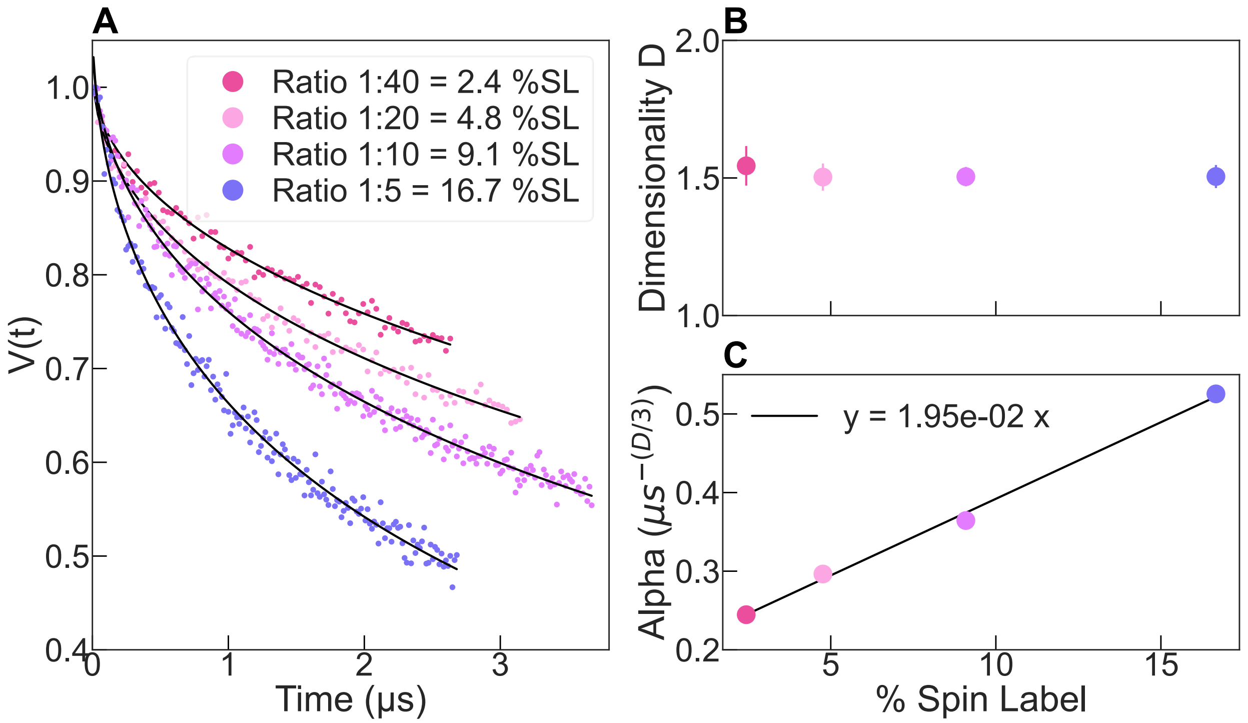


Figure S4: (A) Dipolar evolution time measured for different ratio of labelled to unlabeled protein, effectively changing the PC density along the filament axis. Data are fitted with Equation (1) (black line) to extract (B) the dimensionality and the (C) $\alpha$ parameters.

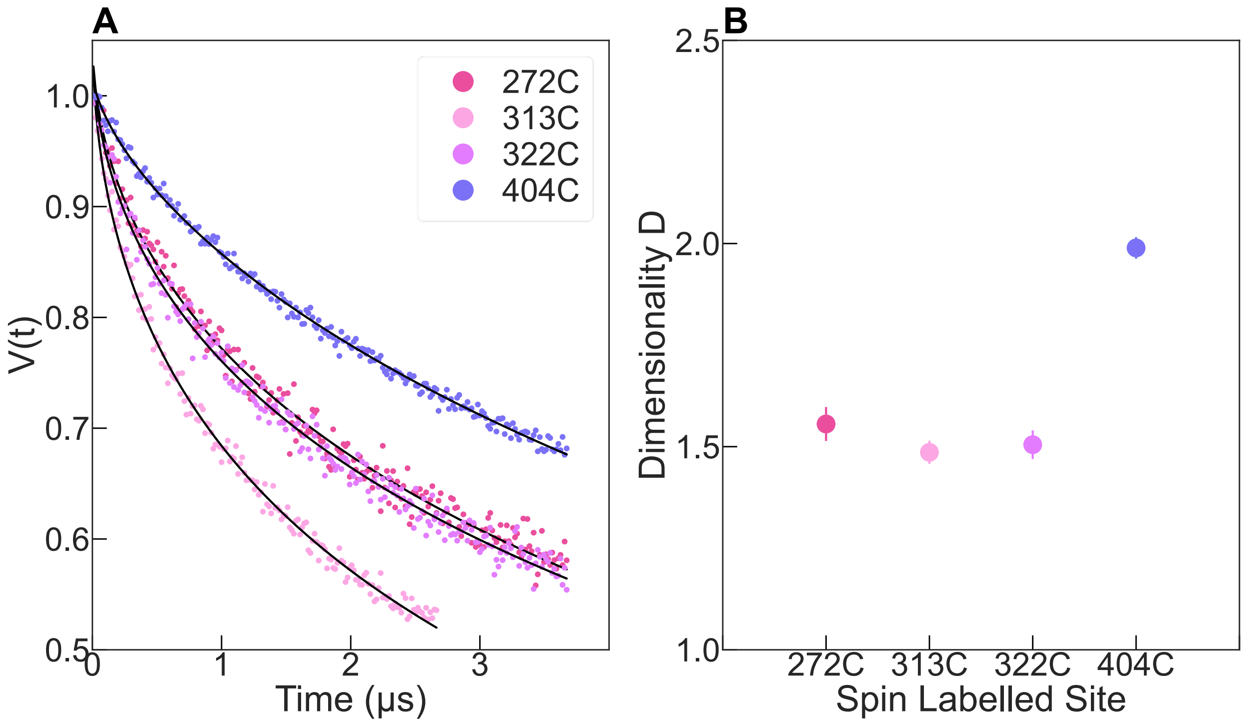


Figure S5: (A) Dipolar evolution time measured in amyloid filaments made with tau protein labelled at different positions, for a SL-tau:noSL-tau ration of 1:10. Data are fitted with Equation (1) (black line) to extract (B) the dimensionality parameter.

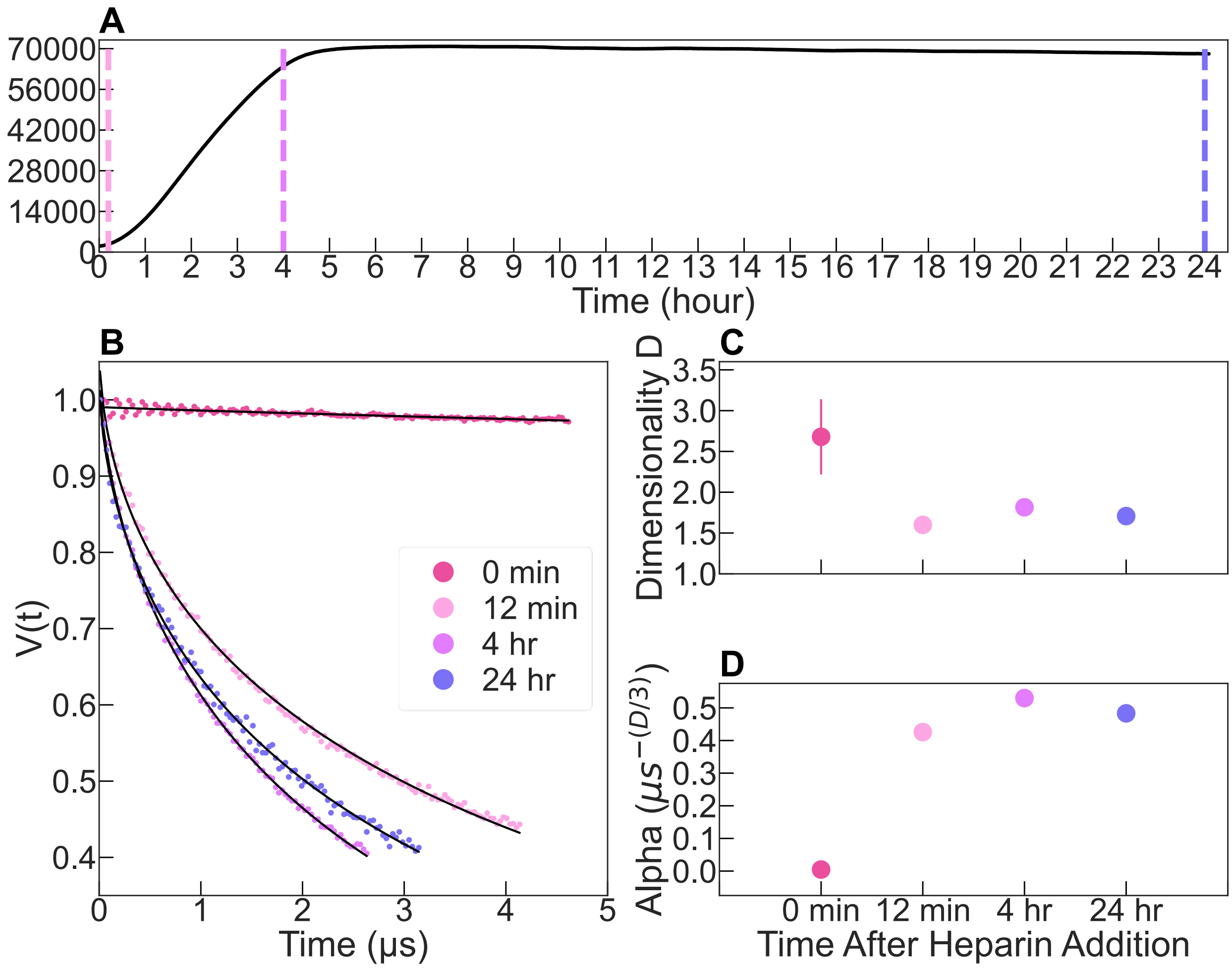


Figure S6: (A) tau187 aggregation was followed by ThT aggregation and samples were flash-frozen before, at 12 min, 4h and 24h after adding heparin (dash lines). (B) ESE decays were fitted with Equation (1) to extract (C) $D$ and (D) $\alpha$ parameters.

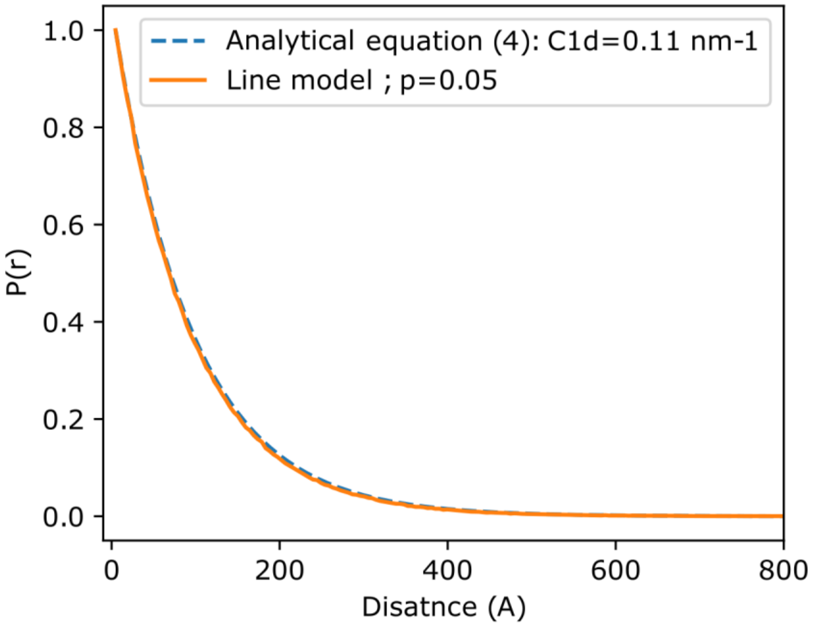


Figure S7: Analytical model from Equation (4) gives the same distribution as the numerical simulation of a line populated with PC (Figure 4 main manuscript).

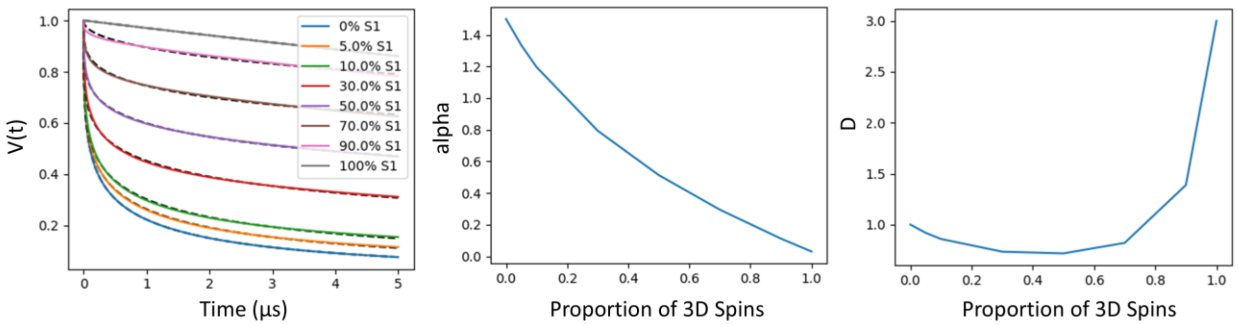


Figure S8: A linear combination of ESE decay from Equation (1) between $D$= 1 and $D$ = 3 (alpha = 0.05) was calculated (left panel). S1 corresponds to the component with $D$=1. This aims at roughly modeling a mixture of spins held together in a 1D matrix and in a 3D matrix (representing an amyloid fiber in a solution containing soluble spin labels). The linear combination of both components was then fitted with Equation (1) and the fitting parameters alpha and $D$ are shown in the central and right panel, respectively.

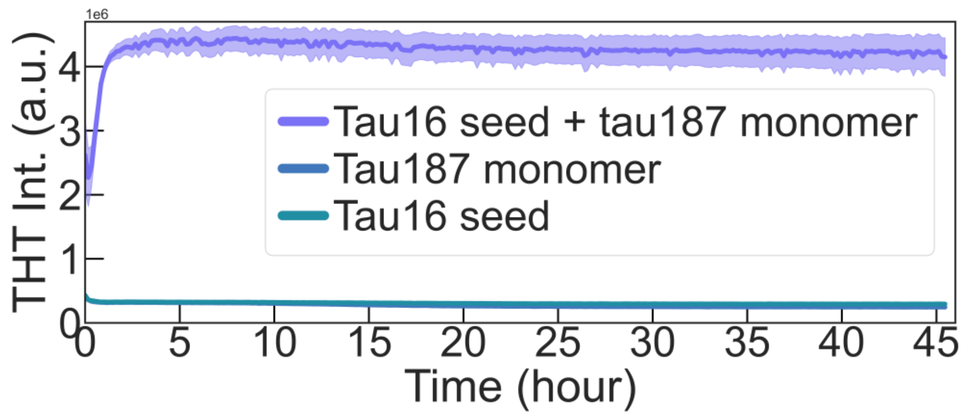


Figure S9: Tau16 seeds induced tau187 aggregation as shown by ThT fluorescence.

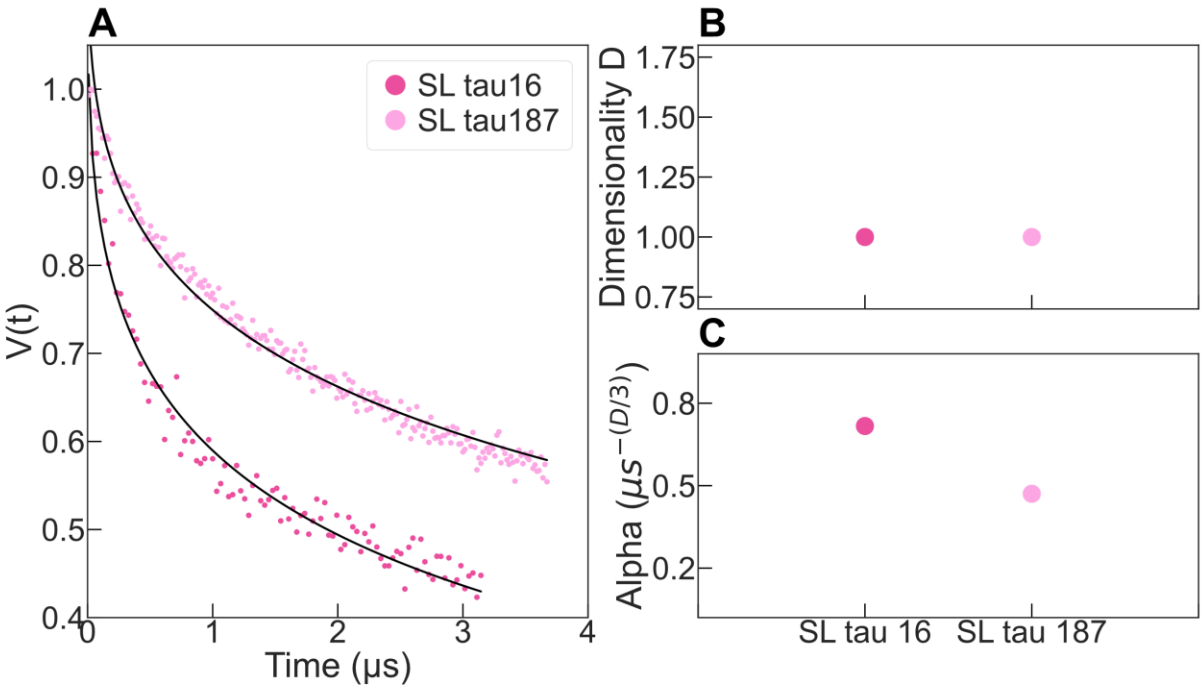


Figure S10. (A) ESE decay for tau16 and tau187 fibers, fitted with Equation (1) with $D$ constrained to 1 (B) and alpha is free (C). Similarly to the conclusion reached from Figure 5 (where $D$ was not constrained), alpha is higher for tau16, revealing a higher spin density.
